# Development of a Prediction Model for Incident Atrial Fibrillation using Machine Learning Applied to Harmonized Electronic Health Record Data

**DOI:** 10.1101/520866

**Authors:** Premanand Tiwari, Katie Colborn, Derek E. Smith, Fuyong Xing, Debashis Ghosh, Michael A. Rosenberg

## Abstract

Atrial fibrillation (AF) is the most common sustained cardiac arrhythmia, whose early detection could lead to significant improvements in outcomes through appropriate prescription of anticoagulation. Although a variety of methods exist for screening for AF, there is general agreement that a targeted approach would be preferred. Implicit within this approach is the need for an efficient method for identification of patients at risk. In this investigation, we examined the strengths and weaknesses of an approach based on application of machine-learning algorithms to electronic health record (EHR) data that has been harmonized to the Observational Medical Outcomes Partnership (OMOP) common data model. We examined data from a total of 2.3M individuals, of whom 1.16% developed incident AF over designated 6-month time intervals. We examined and compared several approaches for data reduction, sample balancing (re-sampling) and predictive modeling using cross-validation for hyperparameter selection, and out-of-sample testing for validation. Although no approach provided outstanding classification accuracy, we found that the optimal approach for prediction of 6-month incident AF used a random forest classifier, raw features (no data reduction), and synthetic minority oversampling technique (SMOTE) resampling (F_1_ statistic 0.12, AUC 0.65). This model performed better than a predictive model based only on known AF risk factors, and highlighted the importance of using resampling methods to optimize ML approaches to imbalanced data as exists in EHRs. Further studies using EHR data in other medical systems are needed to validate the clinical applicability of these findings.

## Introduction

Atrial fibrillation (AF) is the most common sustained cardiac arrhythmia, and its prevalence is increasing^1^; ~5.1M individuals had AF in 2010, and an increase of 9.3-12.1M is anticipated by 2030^2^. Importantly, the increased risk of mortality with AF is almost entirely due to an increased risk of thromboembolic stroke^3, 4^. This risk could be reduced if a moderate or high-risk patient with AF is started on oral anticoagulation^5–17^. A major challenge in the management of patients with AF is that often stroke is the first presentation of AF^18^, indicating that simply waiting for a patient to develop AF may not be the optimal approach to limit the risk of stroke. On the other hand, population-wide screening for AF is not currently recommended ^19–21^, although some suggest that targeted screening may be useful^21^. A model that could predict risk of AF over a 6-month period could be applied to target screening to identify a patient with AF prior to the next clinic visit.

The promise of electronic health record (EHR) data has included the potential to leverage ‘big data’ analytical approaches to predict clinical outcomes within a real-world context. However, despite widespread adoption of EHRs as mandated under the Affordable Care Act^22^, there are limited examples of practical application of EHR data to predict a meaningful clinical outcome^23–27^. In addition to technical limitations of working with data at the scale of the EHR, there are also challenges in performance of external validation across healthcare systems^28–30^. Nonetheless, with increasing availability of cloud-computing^25^ platforms and data storage ^31,32^, as well as scalable computational models that can be developed and potentially shared across healthcare systems, opportunities to apply EHR data to clinical decision making are emerging.

A great deal of enthusiasm has accompanied applications of deep learning^33^ and artificial intelligence to outperform humans in image recognition^34, 35^, text recognition^36, 37^, and games^38^, such as checkers^39^ or Go^40^. However, within the healthcare setting, the ‘black box’ characteristic of machine learning (ML) has caused hesitancy in application. In certain situations, ML approaches, such as support vector machines^41^ or random forests^42^, have been found to produce greater predictive performance than standard regression models^43–45^. More recently, there has been increased recognition that deep-learning models^33, 46^, composed of multiple hidden layers of a neural network rather than a single layer, are better equipped to handle the large amount of data that exists in EHRs. However, in order to understand how these approaches can be applied to a clinical situation, such as prediction of incident AF, additional study is needed.

In this investigation, we developed and tested an ML model to predict 6-month incidence of AF using EHR data. We conducted a systematic examination of EHR data sampled from 2.3 million individuals, in whom we have harmonized 26,000 features, including diagnostic codes and medications under the OMOP common data model. Among the characteristics we examined in this developmental process includes the appropriate use of data reduction techniques, data resampling to manage dataset imbalance, and identification of a classification algorithm based on training time and accuracy.

## Methods

### Study Population and Case Ascertainment

The UCHealth hospital system includes 3 large regional centers (North, Central, South) over the front range of Colorado that share a single Epic instance, which allows data from all centers to be pooled into a single data warehouse, a copy of which is located on the Google cloud platform. This warehouse of data was queried using Google BigQuery to create a dataset and conduct analyses directly on the Google cloud platform, where an array of machine-learning tools can be run on virtual machines. To create our study dataset, we applied a classification approach based on predicting risk of incident AF over a 6-month period, as this timeframe is also the standard follow-up time for most cardiovascular providers. We performed a SQL query on the UCHealth EHR for subjects with new diagnosis of AF obtained over a 6-month interval. To identify cases, we filtered out all patients with prevalent AF on first encounter, and then over 6-month intervals (from each encounter), assigned patients to a ‘case’ classification if they had AF diagnosed by ICD code (ICD-9 427.31 or ICD-10 I48.91) within that interval. Once a patient was designated a case, he/she was removed from the pool, and all non-case patients without AF were designated as ‘controls’. Data was available in the EHR for the period from January 1, 2011 until October 1, 2018 for 2.3M subjects. This study protocol was approved for analysis of de-identified data (limited dataset with dates included) by the University of Colorado Institutional Review Board.

### Common Data Model and Data splitting

To provide the opportunity for validation of findings in this study, we used a common data model for EHR data, based on the Observational Health Data Sciences and Informatics (OHDSI) collaboration, which uses the Observation Medical Outcomes Partnership common data model (OMOP-CDM)^47^. The OMOP CDM is a mapping of the raw EHR data to a harmonized dataset; for this investigation, we used this CDM with 26k variables (i.e., features) from the EHR, including diagnosis codes and medications. These values are time-stamped with the time of entry into the medical record, which is used to correlate with the timing of the outcome of interest. Features are encoded using one-hot encoding, and were collected cumulatively from the time of first encounter until diagnosis of AF (cases) or end of follow-up (controls). To reduce the time for computation, as well as preserve additional data for future validation studies within our medical system, we obtained a random sample of 412,291 subjects (407,550 controls and 4741 cases), which was then split into training (80%) and testing (20%) sets to compare the models developed in this investigation. The training set underwent an additional split to create a validation set (10% of the training set) for comparing unsupervised stacked autoencoder models (see below). Figure 1 displays the data management scheme for this investigation.

**Figure 1.**
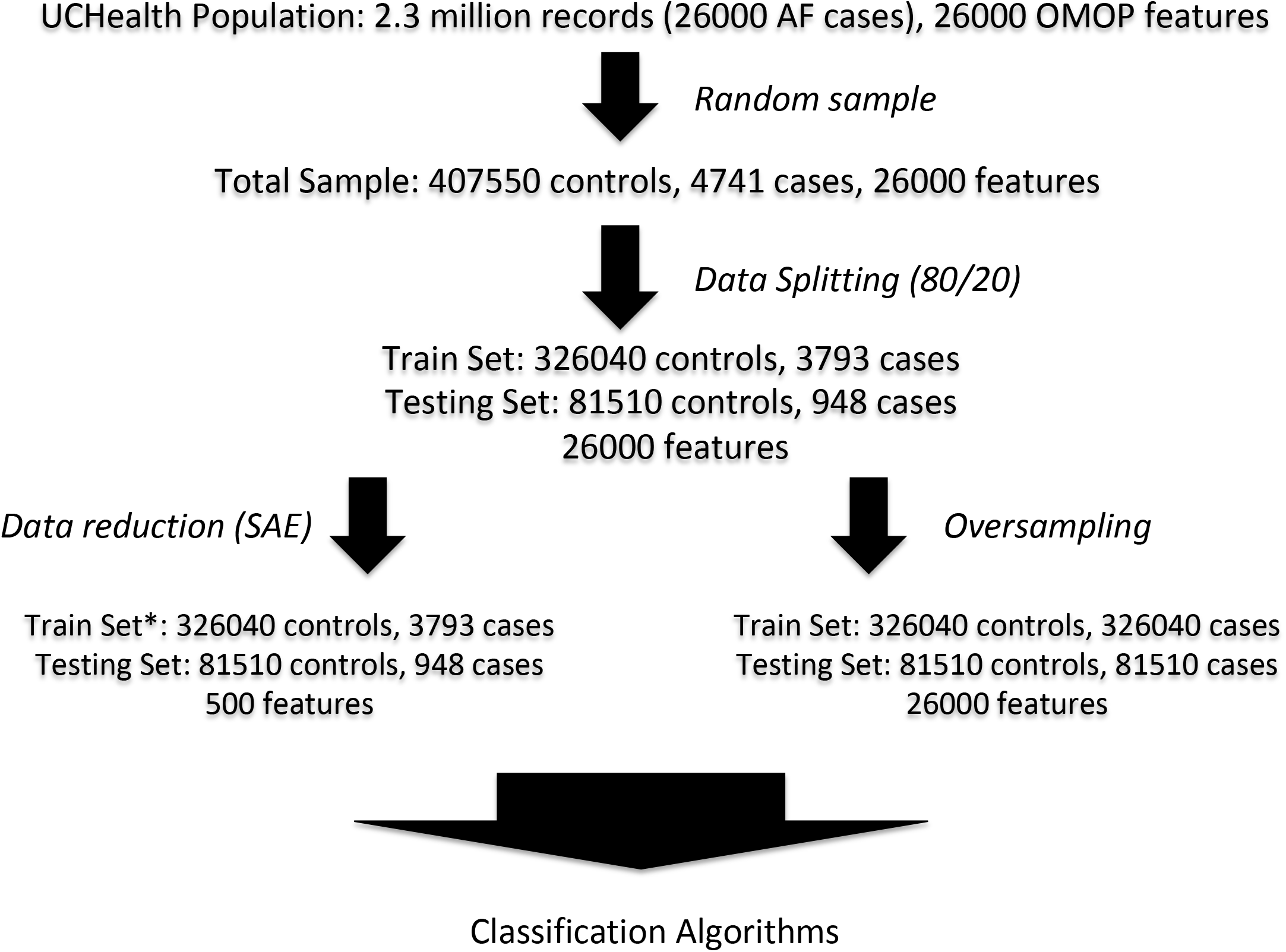
Study sampling design. *Note that for comparison of stacked autoencoders (unsupervised learning), the training set was split into an additional validation set (10% of training set). See *Methods* for details.

### Model Development

Hyperparameter tuning was performed using iterative random sampling of 10,000 records for manual grid search (neural networks), and 10-fold cross validation for automated grid search (for other machine learning approaches). See *Supplemental Methods* for details. Unsupervised analysis was performed first using principal component analysis to examine overall data structure. For dimensionality reduction, we employed stacked autoencoders^48^ using fully connected neural networks of several architectures, with a goal to identify the lowest replication error (cross-entropy loss) within the validation set (10% of training set). We also conducted analyses using the full (non-reduced) feature set of 26k features for comparison.

We examined several strategies for resampling, including random oversampling, SMOTE^49^, random undersampling, and cluster centroid. To identify the best resampling approach, we used random forest classifier, as pilot analyses using a smaller dataset suggested this approach might be superior to other ML approaches. We also compared with a model using no resampling (imbalanced).

Once we identified an optimal resampling approach, we compared several classification algorithms, including naïve Bayesian classification, random forest classification, boosted gradient classification, support vector machines, one-layer fully connected neural networks (shallow) and multiple layer fully connected neural networks (deep). Model comparison was based on area-under-curve and F_1_ statistic^50–52^. Computation time includes all prior data sampling and algorithm performance. Once an optimal model and resampling approach were identified, we conducted sensitivity analysis using several alternative resampling and modeling approaches in combination to ensure that the combination (dimensionality reduction, resampling, and classification algorithm) identified was indeed optimal. Precision-recall and receiver-operator characteristic curves, as well as feature importance plots, were created for the optimal model for manual inspection.

### Validation of Developed Model

The optimal model was then compared with an unregularized logistic regression model based on presence of known clinical predictors of AF, including hypertension (ICD-9 and 10–Hypertension: 401.x and I10, Obesity: 278.x and E66.9, Diabetes: 250.x and E 11.9, Coronary disease: 414.x and I25.1x, Heart failure: 428.x and I50.9, Valvular heart disease: 424.x and I08).

### Computation and Analysis

All analyses were run on Google Cloud Platform, using 96 CPUs and 620 GB of RAM. Scripts were composed in Python (version 3) and were run on Jupyter Notebook with Tensorflow platform on the Google Cloud Platform. Machine learning packages included *scikit-learn* and *keras.* Confidence intervals were calculated using Wald method^53, 54^, although almost all were within the rounding error of the estimates due to the large testing sample size (N = 82,458), and are not displayed. See *Supplemental Methods* for additional details.

## Results

Across the entire UCHealth population of 2.3M, we identified over 26k patients with 6-month incident AF (Table 1). Although essential hypertension was the most common data element in both groups, patients with 6-month incident AF had more standard cardiovascular diagnoses compared to the larger population who were never diagnosed with AF. From this population, a random sample selected for model development included 407,550 controls and 4,741 cases, which were split into a training set (80%) of 326,040 controls and 3,793 cases and a testing set (20%) of 81,510 controls and 948 cases (Figure 1).

**Table 1.** Description of UCHealth cohort by AF diagnosis. Provided are the mean age and gender by 6 month AF diagnosis, as well as the top 10 diagnosis codes, according to whether AF was diagnosed within a 6-month period.

|  | **No AF** | **6-month Incident AF** |
| --- | --- | --- |
| **Number (%)** | 2.3M (98.85%) | 26K (1.15%) |
| **Age (Mean ± SD)** | 42.48 ± 22.28 | 71.14 ± 16.28 |
| **Female sex (%)** | 54.79% | 45.2% |
| **#1 code** | 320128 Essential hypertension<br>15.6%<br>SNOMED:59621000 | 320128 Essential hypertension<br>44.66%<br>SNOMED:59621000 |
| **#2 code** | 254761 Cough<br>10.03%<br>SNOMED:49727002 | 432867 Hyperlipidemia<br>32.42%<br>SNOMED:55822004 |
| **#3 code** | 200219 Abdominal pain<br>9.89%<br>SNOMED:21522001 | 77670 Chest pain<br>17.08%<br>SNOMED:29857009 |
| **#4 code** | 432867 Hyperlipidemia<br>9.23%<br>SNOMED:55822004 | 318800 Gastroesophageal reflux disease<br>14.94%<br>SNOMED:235595009 |
| **#5 code** | 257011 Acute upper respiratory infection<br>8.87%<br>SNOMED:54398005 | 312437 Dyspnea<br>14.91%<br>SNOMED:267036007 |
| **#6 code** | 77670 Chest pain<br>8.86%<br>SNOMED:29857009 | 201826 Type 2 diabetes mellitus<br>14.65%<br>SNOMED:44054006 |
| **#7 code** | 25297 Acute pharyngitis<br>8.15%<br>SNOMED:363746003 | 42872402 Coronary arteriosclerosis in native artery<br>13.38%<br>SNOMED:1641000119107 |
| **#8 code** | 378253 Headache<br>7.67%<br>SNOMED:25064002 | 40481919 Coronary atherosclerosis<br>13.35%<br>SNOMED:443502000 |
| **#9 code** | 442077 Anxiety disorder<br>7.66%<br>SNOMED:197480006 | 439926 Malaise and fatigue<br>12.80%<br>SNOMED:271795006 |
| **#10 code** | 436096 Chronic pain<br>7.39%<br>SNOMED:82423001 | 254761 Cough<br>12.38%<br>SNOMED:49727002 |

To explore the data structure of the EHR data prior to modeling, we performed principal component analysis (PCA), where we noted a large amount of overlap between components in patients with and without 6-month incident AF within the first two principal components (Figure 2A). We noted that only a small amount of overall variability was explained by the first few components, and that there was not a clear plateau present over the first 500 components that were analyzed (Figure 2B). We then created several architectures of stacked autoencoder (SAE) neural networks, using regularization techniques such as drop-out and several activation functions. We found that a deep neural network [10000, 2000, 500, 2000, 10000] with three encoding and three decoding layers, dropout, and sigmoid activation function, resulted in the lowest reconstruction (validation) error (Table 2).

**Figure 2.**
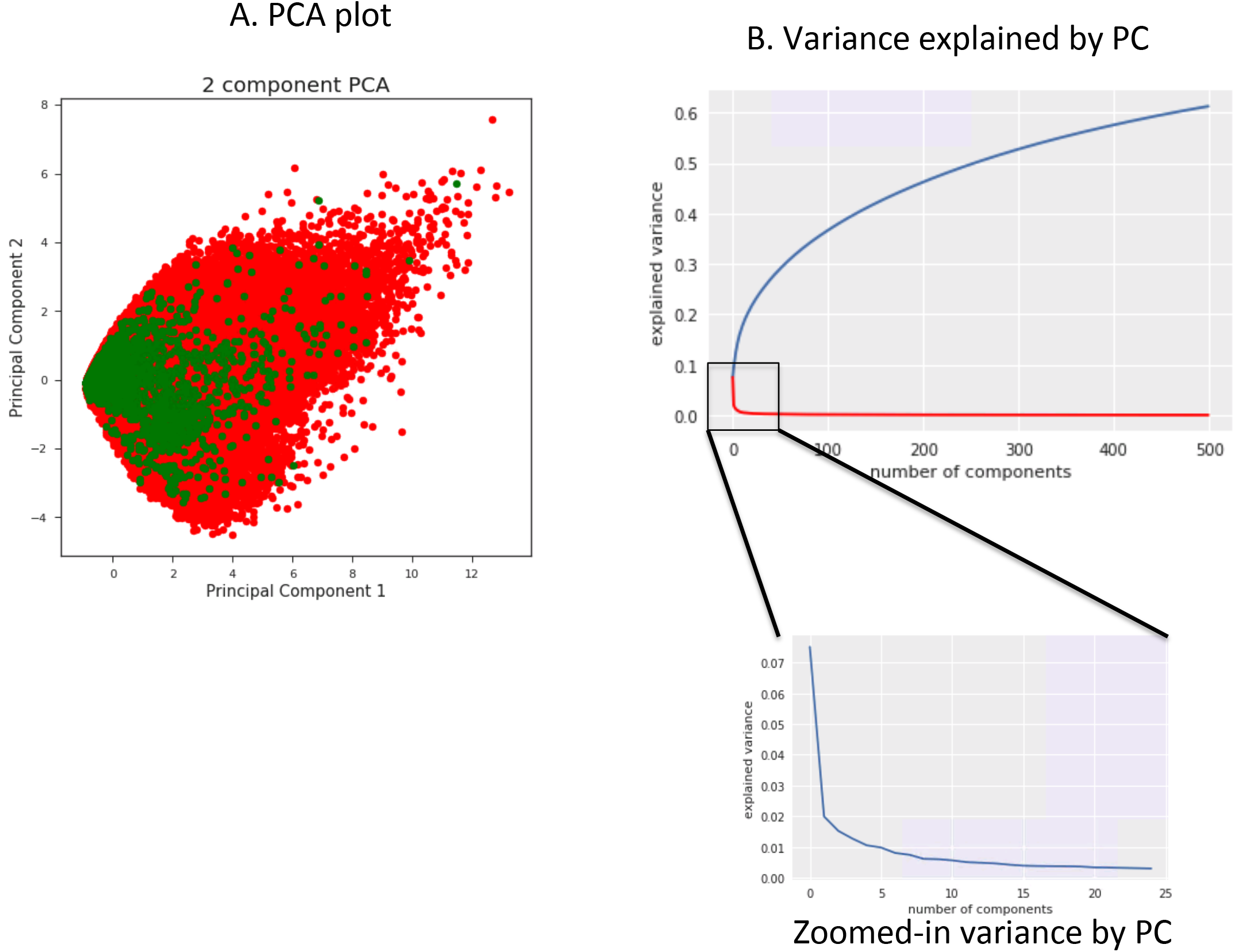
A. First two principal components of codes (color labeled by case and control). Red = control, green = case. **B. Variance explained by PC.**

**Table 2.** Comparison of Stacked Autoencoders.

| **Architecture of Stacked Autoencoders** | **Validation Error** |
| --- | --- |
| 26000-13000-6000-2000-500-2000-6000-13000-26000 | 0.035 |
| 26000-13000-6000-2000-500-2000-6000-13000-26000, dropout | 0.049 |
| 26000-10000-500-10000-26000,dropout | 0.038 |
| 26000-10000-2000-500-2000-10000-26000, dropout, sigmoid activation | 0.007 |
| 26000-500-26000 | 0.067 |
| 26000-10000-500-10000-26000, dropout, sigmoid activation | 0.045 |
| 26000-10000-2000-100-2000-10000-26000, dropout, sigmoid activation | 0.019 |

We then examined the role of undersampling and oversampling methods to identify the optimal approach to manage the imbalance between cases and controls that we identified in this dataset. Using a random forest classification algorithm with three-layer sigmoid/dropout SAE (see above), we found that SMOTE not only provided the shortest training times, but it also resulted in the best classification F_1_ score (Table 3), compared with other methods.

**Table 3.**
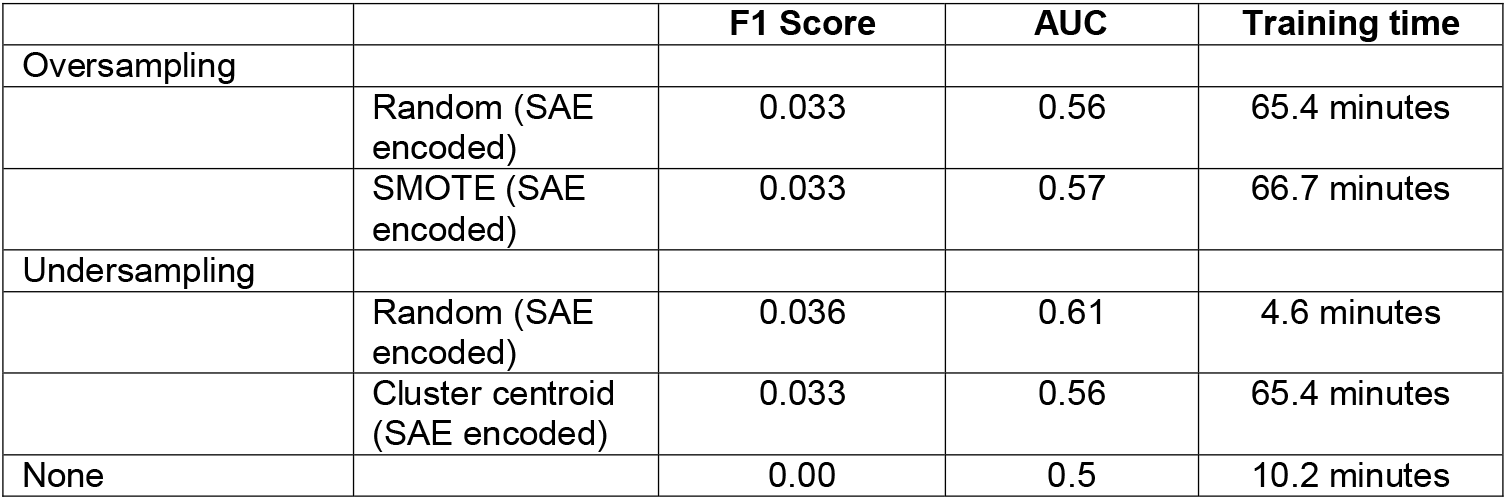
Comparison of Resampling Strategies. Sampling comparison from Random Forest model; performed using SAE encoded as specified.

|  |  | **F1 Score** | **AUC** | **Training time** |
| --- | --- | --- | --- | --- |
| Oversampling |  |  |  |  |
|  | Random (SAE encoded) | 0.033 | 0.56 | 65.4 minutes |
|  | SMOTE (SAE encoded) | 0.033 | 0.57 | 66.7 minutes |
| Undersampling |  |  |  |  |
|  | Random (SAE encoded) | 0.036 | 0.61 | 4.6 minutes |
|  | Cluster centroid (SAE encoded) | 0.033 | 0.56 | 65.4 minutes |
| None |  | 0.00 | 0.5 | 10.2 minutes |

Using the SMOTE resampling strategy combined with three-layer SAE (dropout + sigmoid activation), we then examined several classification algorithms to identify a potential ‘overall’ best model. Several models, specifically support vector machines, did not converge after over 24 hours of processing and were not included. Among the approaches examined, we found that random forest classification was superior to other methods that included regularized regression, boosted gradient descent, shallow, and deep neural networks (Table 4).

**Table 4.**
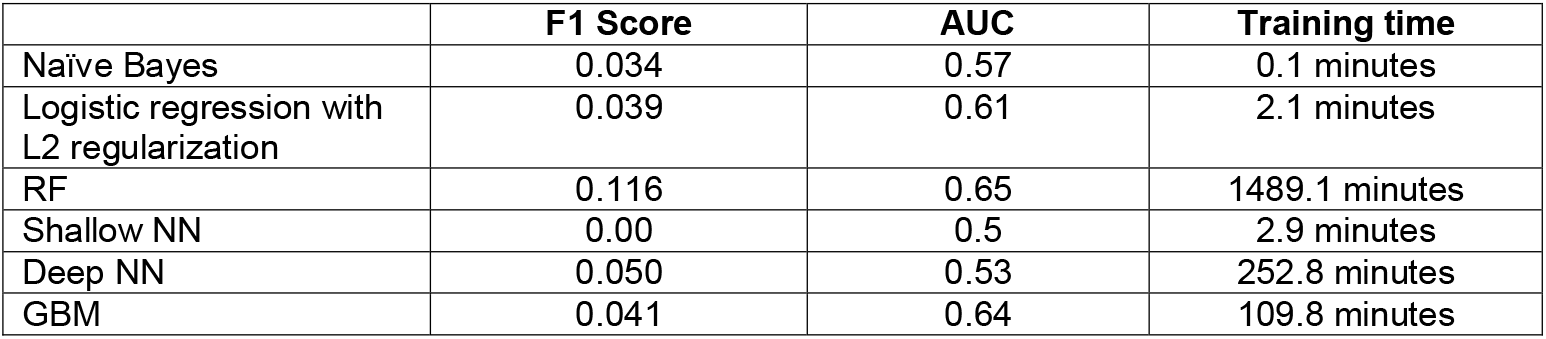
Comparison of machine learning approaches. Using SMOTE resampling technique and all features. F1 and AUC calculated from model applied to held-out testing set (20%); training time is for training of training set (80%)

To ensure that our modeling combination did not favor a particular combination of dimensionality reduction, resampling, and classification algorithm, we then performed a sensitivity analysis using various combinations of each. We found that a model that used the raw features as input (no dimensionality reduction), SMOTE and random forest classifier (F1 0.12, AUC 0.65) performed better than other combinations, including raw features, SMOTE and 2-layer neural network (F1 0.05, AUC 0.53), and was overall the best predictive model identified from our EHR. This model had a specificity of 94.7%, sensitivity of 35.74%, negative predictive value of 99.3% and positive predictive value of 6.93% at a probability (decision) cutoff of 0.5. The confusion matrix for the model is displayed in Supplemental Figure 1. As shown in Figure 3A, several additional probability cutoffs for classification before 0.4 provided over 95% sensitivity, at the expense of a significant decrease in specificity. Feature importance was examined for the optimal model, which found that none of the features most common across the population (Table 1) were of high importance in classification of 6-month AF (Supplemental Table 1 and Supplemental Figure 2), although several features predominant in incident AF patients had non-zero importance (42872402 Coronary arteriosclerosis in native artery: 1.61e-05; 254761 Cough: 4.11e-06; 201826 Type 2 diabetes mellitus: 1.96e-07). Calibration curves also revealed relatively poor classification across all probability thresholds (Supplemental Figure 3) for several models.

**Figure 3.**
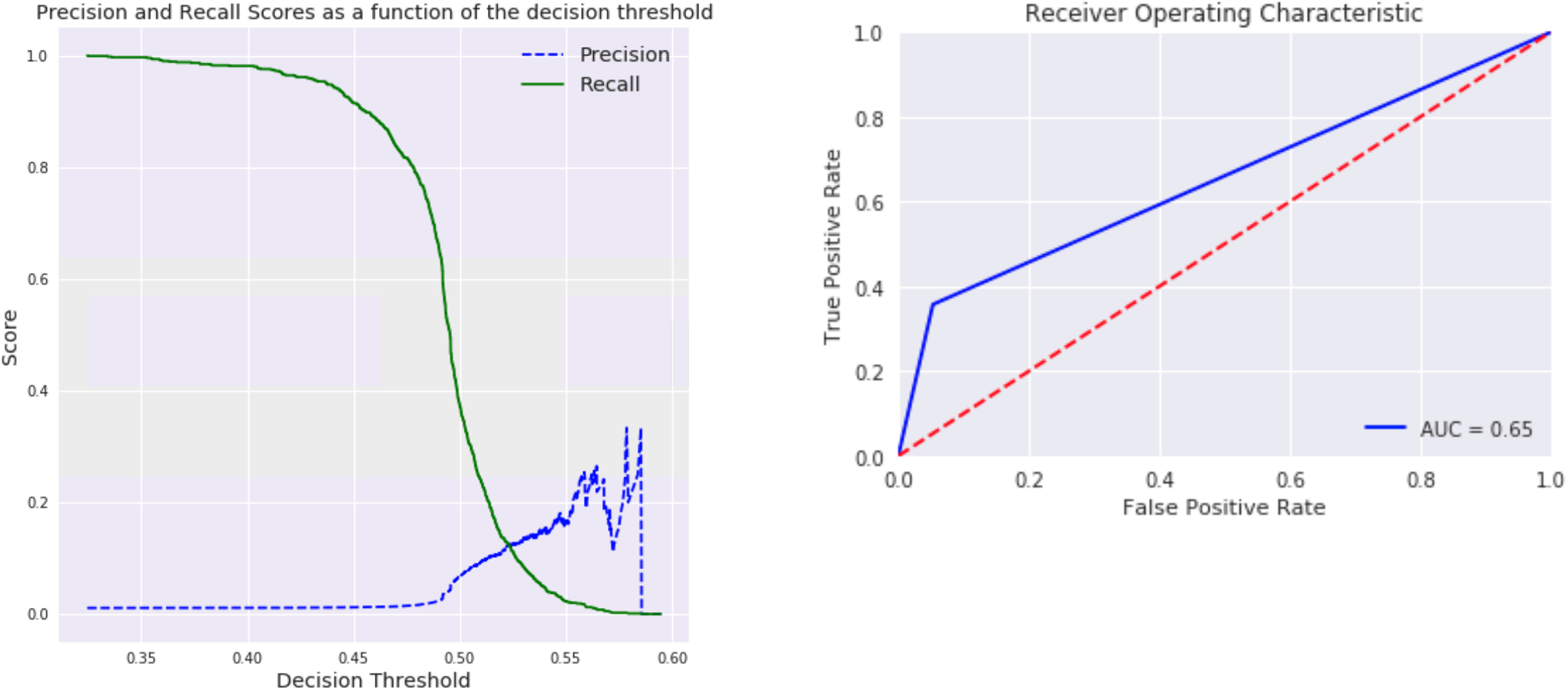
A. Precision-recall curve for optimal model. B. ROC curve for optimal model.

Finally, we found that the optimal model (raw features, SMOTE, random forest classifier) performed better than an unregularized logistic regression model based on known AF risk factors, which had an F_1_ score of 0.073, AUC of 0.64 with SMOTE resampling prior to fitting (training), and an F_1_ score of 0.00 and AUC of 0.5 without resampling prior to fitting.

## Discussion

In this investigation of development of a machine-learning model using harmonized EHR data for predicting 6-month incident AF, we found that a random forest classifier created from raw feature inputs with SMOTE oversampling provided better classification than any other combinations of approaches tested, including deep learning models and known risk factors of AF. These results are significant because in addition to motivating future investigations to apply ML methods to EHR data to identify patients at risk for AF, they also incorporated harmonized data (OMOP-CDM), which means that the optimal model can not only be directly applied in our EHR, but to data from the EHR of any other medical institution participating in OMOP/OHDSI. In clinical application, our model could thus be inserted directly back into the user interface to guide targeted screening patients at risk of AF, including development of prospective follow-up studies to use the prediction for targeted screening for AF, including routine ECGs, implantable, or wearable devices.

However, there are several reasons for hesitancy before taking these results directly back to the bedside to guide clinical management, without additional investigation. First, the model that we identified was not extremely accurate, with an F1 score well under 20%, and a sensitivity of 35.74% based on a cutoff probability of 0.5 for risk. Although we noted that the probability threshold can be lowered to improve sensitivity of classification, the drop in specificity, and number of false-positives, with such an approach would result in a large number of patients undergoing unfruitful screening. Second, we found that the features identified as important in the classification process were atypical, with many of unclear association with AF based on known pathophysiology^55, 56^ and clinical risk factors^57, 58^. Although this finding may reflect the fact that none of the features was particularly strong in predicting risk of AF, as evident from the low feature-importance scores, it also suggests that the model may be overfitting our population (despite validation in a held-out testing set). Importantly, because we performed a harmonization step prior to the modeling process, the necessary external population validation to explore the possibility of overfitting is simply a matter applying this model directly to OMOP-CDM from an outside EHR. It is also important to keep our findings in context as there is a great deal unknown about clinical risk in development of AF, and one need look no further than to the field of genetic investigation, where whole-genome approaches have identified novel risk loci whose role in AF pathophysiology remains poorly understood^59, 60^, but which provide far superior prediction over many candidate genes.

In the process of developing of a 6-month risk prediction model for AF, we made several important observations about the application of machine learning to EHR data. First, we found that dimensionality reduction was inferior to use of raw feature inputs for predicting incident AF. This finding likely reflects the sparsity of EHR data, which resulted in a significant loss of information even using dimensionality reduction methods with very low reconstruction loss. This finding implies that within our population, the strongest predictors of 6-month incident AF are relatively rare, and thus minimized with dimensionality reduction approaches. The relatively obscure features identified in the feature importance evaluation supports this contention. To our knowledge, this phenomenon has not been described in EHR-analysis approaches, many of which apply some form of dimensionality reduction^61^, although further exploration is likely necessary.

Second, we found that for a rare condition like 6-month incident AF (1.2% of the total population), oversampling to rebalance the data was superior to using the imbalanced dataset and undersampling, regardless of the classification algorithm applied. In fact, the majority of the ML approaches tested failed to provide any added improvement (F1 0.0 and AUC 0.5) without sample rebalancing. This finding is likely reflective of the increase in power obtained with oversampling, although further work is needed to understand why this particular approach was superior.

Finally, we found that random forest classification provided the optimal classification algorithm, with better classification than a deep neural network approach when the input was raw features. This finding demonstrates that although deep learning approaches may be superior for classification of structured datasets, such as in image^62^ or voice recognition^63^, they are not always optimal over other ‘standard’ ML algorithms, and highlights the importance of examining all approaches for each classification problem, rather than assuming a giving approach is optimal.

### Strengths

In addition to the insights above, there are several additional strengths noteworthy in this investigation. First, all models were created using a harmonization scheme (OMOP common data model) that could allow for direct application and validation to data mapped from a separate EHR. Such harmonization allows for the opportunity to explore transfer-learning^64^ approaches, which could provide additional insight into similar and divergent AF risk factors across populations. Second, we conducted a systematic approach to identify the best dimensionality reduction, resampling, and classification algorithm for this outcome. Further work in other outcomes is needed to determine if the combination we identified for predicting 6-month incident AF is also optimal for prevalent or longer-term AF prediction, as well as for outcomes that are more or less common than AF. Finally, we examined a dataset of over 2 million subjects, which provided more than enough sample size from our single institution to conduct cross-validation and out-of-sample validation. This power from use of big data is possible by the unique circumstances of our relationship with Google Cloud Platform, although many other EHRs are moving to the cloud, providing further opportunities for development and testing.

### Limitations

There were several limitations in this study, many of which are the subject of future, more targeted investigations. For one, our study included a very simple method for the temporal relationships between features in our dataset, which did not account for time-varying effects or censoring. An AF event that occurs the day after an encounter is modeled the same as one occurring the day before a subsequent encounter, and a diagnosis or medication that was given one month before the AF diagnosis was weighted the same as one given 4 years prior. While we suggest that the approach we employed for this investigation is reasonable based on the typical 6-month follow-up schedule for patients seen in cardiology clinic, we realize that additional information about temporal risk will be needed for more accurate prediction approaches. More sophisticated methods, such as recurrent neural networks^61, 65^ or parametric survival functions^66^ could provide more accurate prediction in future investigations. A second weakness is that we excluded some additional data elements, such as lab values and diagnostic test results, which may have had prognostic value for predicting 6-month AF^67, 68^. Some of these values have been difficult to harmonize across datasets via OMOP-CDM, and others suffer from high variability in inter-institutional measurement, such as echo measures of diastolic function^57, 69^. Nonetheless, there are many additional biomarkers^67, 68^ likely to have a more ‘biological’ relationship to risk of AF than a diagnostic code, and future applications that include this information would be expected to provide both predictive and inferential knowledge about risk of AF. Finally, although the systematic, harmonized approach we employed in this study holds potential for cross-institutional validation, much work is needed in terms of data sharing before actual testing can be performed. Our group and others are working in this direction, and the hope is that sometime in the not-to-distant future, all EHRs will incorporate a ‘standard’ risk prediction model for AF and many other conditions.

In conclusion, we studied the development of an ML model to predict 6-month risk of AF using harmonized EHR data and found that the combination of raw feature inputs, SMOTE oversampling, and random forest classification provided superior prediction than other models, including one with known clinical risk factors. Further work is needed to explore the technical and clinical applications of this approach model to improving outcomes.

## Acknowledgements

This work was funded by grants from the National Institute of Health/NHLBI (MAR: 5K23 HL127296).

