## Supplemental Material for "Development of a Prediction Model for Incident Atrial Fibrillation using Machine Learning Applied to Harmonized Electronic Health Record Data"

**Supplemental Methods**

Hyperparameter Tuning

Grid search was used to identify the optimal hyperparameters for all models, as relevant to the available hyperparameters for each machine-learning approach, with range 3 – 9 hyperparameter values evaluated per search. For algorithms included in the *sklearn* package (see below), the *sklearn.model_selection.GridSearchCV* function was applied with 10-fold cross-validation. For neural networks, iterative random sampling of 10,000 records was performed using the *sklearn.utils.shuffle* package to generate a separate dataset for each value in the grid, and models were fit using each value within the grid. For *keras* package (i.e., neural networks), grid search was performed using different activation functions (sigmoid, tanh, LeakyReLU, PReLU), different numbers of neurons per layer (range 500 – 15000), different number of hidden layers (range 1 – 5), learning rate (range 0.001 – 0.05), dropout (range 0.2 – 0.5), and early stopping (with and without).

Unsupervised Learning

For principal component analysis (PCA), the Python package *sklearn.decomposition.PCA* was used with 500 PCA components. For stacked autoencoders, the *keras.models.Sequential, keras.layers.Dense, keras.layers.Activation, keras.layers.Dropout, keras.layers.advanced_activations.LeakyReLU, and keras.layers.advanced_activations.PReLU.* Models were developed using 90% of the training set, and compared using 10% of the training set (validation set).

Resampling

For resampling comparison, we employed the *imblearn.over_sampling_RandomOverSampler, imblearn.over_sampling_SMOTE, imblearn.under_sampling.RandomUnderSampler, imblearn.under_sampling.ClusterCentroids, and imblearn.over_sampling_SMOTETomek* packages. Of note, SMOTE-Tomek did not achieve convergence after multiple hours of computation, and was excluded from this analysis.

Supervised Analysis

See above for hyperparameter search strategy for each model. L2 Regularized logistic regression was performed using *sklearn.linear_model.LogisticRegression* with inverse regularization strength (C parameter) of 1000, tolerance of 0.0001, and ‘sgd’ optimization algorithm. Naïve Bayesian analysis was performed using *sklearn.naive_bayes* with default settings. Random forest classification was performed using *sklearn.ensemble.RandomForestClassifier* with 200000 estimators (trees), with max depth of 5. Shallow neural network was performed using *Keras* with a single fully connected layer of 5000 neurons, with tanh activation and softmax output, with dropout of 20%, Adam optimization (Learning rate = 0.005, epsilon = $1x10^{-8}$, decay = 0.0) and sparse cross-entropy loss. Deep neural network included two hidden layers with 5000 neurons each, tanh activation, dropout 20%, Adam optimization (Learning rate = 0.005, epsilon = $1x10^{-8}$, decay = 0.0), and sparse cross-entropy loss and early stopping. Gradient boosted classifier was performed using *xgboost.XGBClassifier* and Random Forest classifier, using parameters above. Support vector classification was attempted using *sklearn.svm.SVC* with default hyperparameters, although models failed to converge after many hours of computation, and were not included.

Sensitivity analysis was performed using the random forest and deep (2-layer) neural network, combined with full feature set, optimal encoded (middle) layer from the stacked autoencoder (26000-10000-2000-500-2000-10000-26000, with dropout and sigmoid activation), random undersampling, and SMOTE oversampling. This comparison was performed on the test set (20% of total).

Calibration curves

Calibration curves were created using *sklearn.calibration.calibration_curve* function with 10 bins for the final model, and three additional comparative models.

Supplemental Table 1. Feature Importance of Final model

| **Feature Importance –Final Model** |
|---|
| 4005186<br>Drug measurement<br>121278003 |
| 80593<br>Urinary complication<br>49698005 |
| 4186577<br>Iatrogenic Cushing's disease<br>41299009 |
| 195513<br>Carcinoma in situ of vagina<br>92791005 |
| 19077763<br>Guaifenesin 20 MG/ML Oral Solution<br>310604 |
| 36713918<br>Somatic dysfunction of lumbar region<br>718929000 |
| 45757639<br>Mammographic microcalcification of breast<br>27931000119107 |
| 4006457<br>Trisomy 21- meiotic nondisjunction<br>205615000 |
| 256909<br>Primary pulmonary coccidioidomycosis<br>88036000 |
| 4341231<br>Tracheo-esophageal fistula without atresia of esophagus<br>235640006 |

**Supplemental Figure 1. Confusion Matrix of Final Model**

**Supplemental Figure 2. Feature Importance plot of Final Model**

**Supplemental Figure 3. Calibration curves for final model (Random Forest = RF), Naïve Bayes, and L2 Logistic regression.**

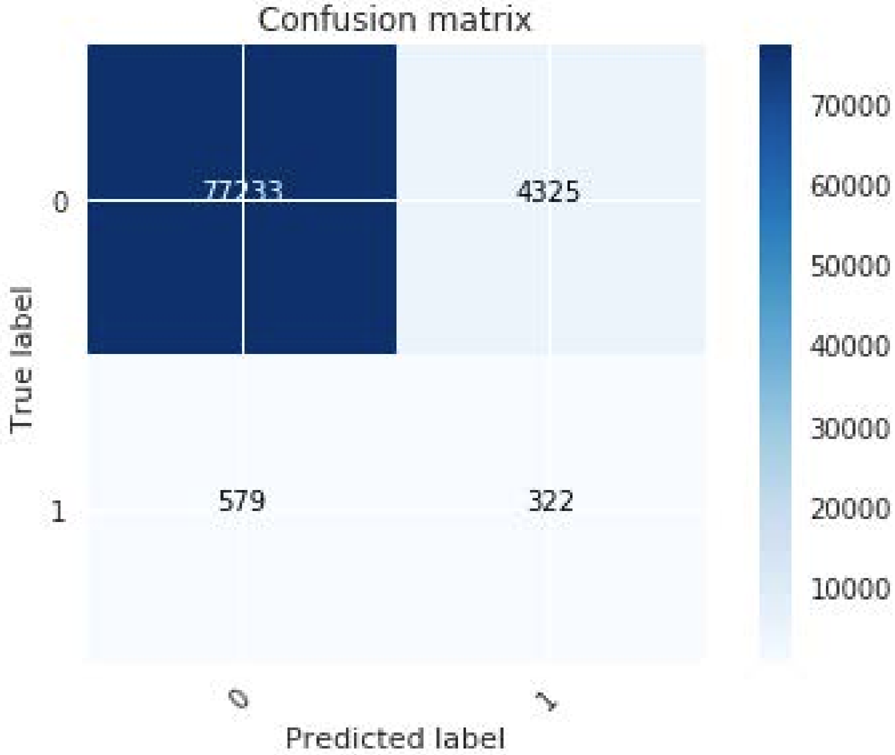
Confusion matrix
77233
4325
579
322
0
1
True label
Predicted label
70000
60000
50000
40000
30000
20000
10000

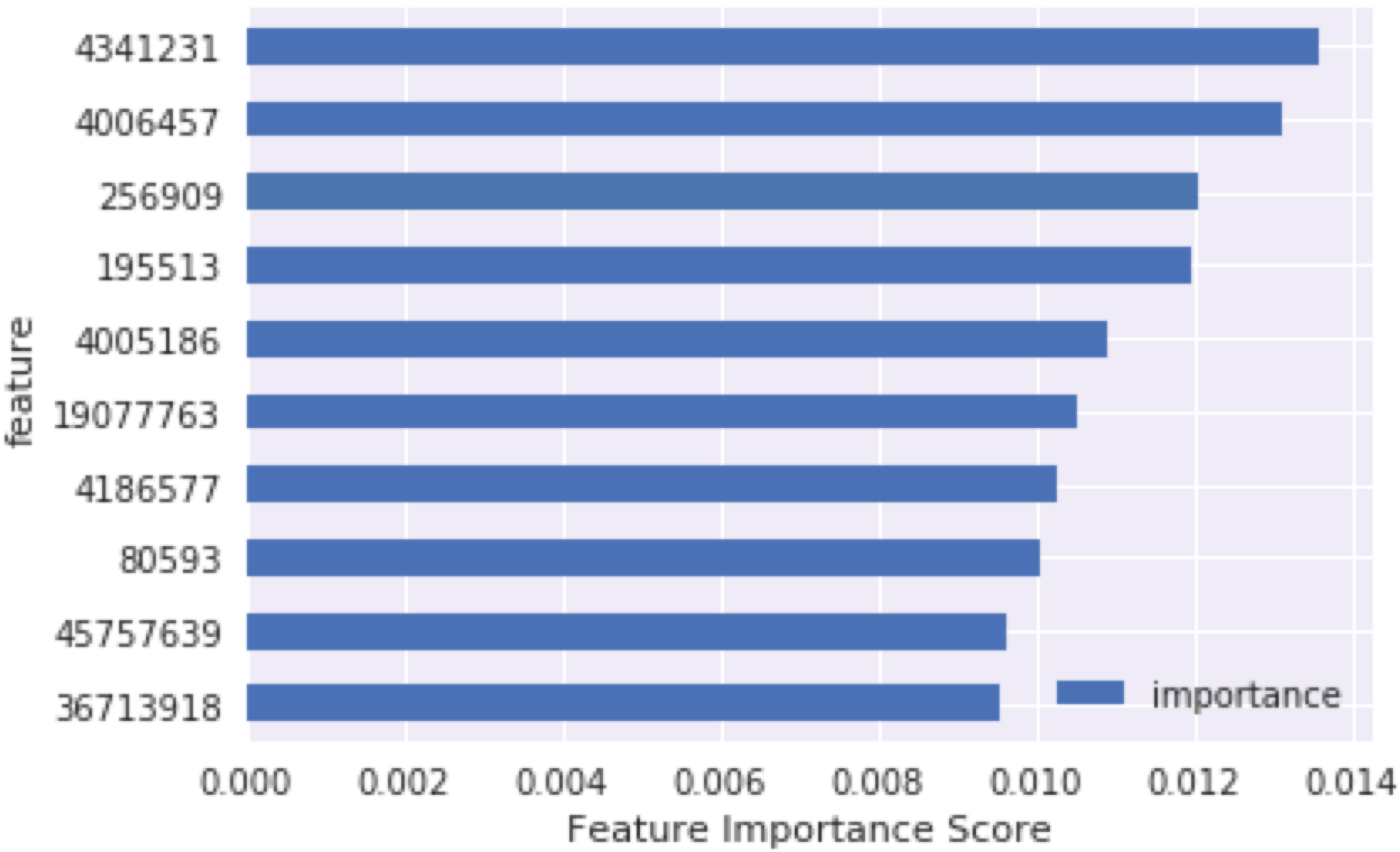

4341231
4006457
256909
195513
4005186
19077763
4186577
80593
45757639
36713918
feature
importance
0.000
0.002
0.004
0.006
0.008
0.010
0.012
0.014
Feature Importance Score

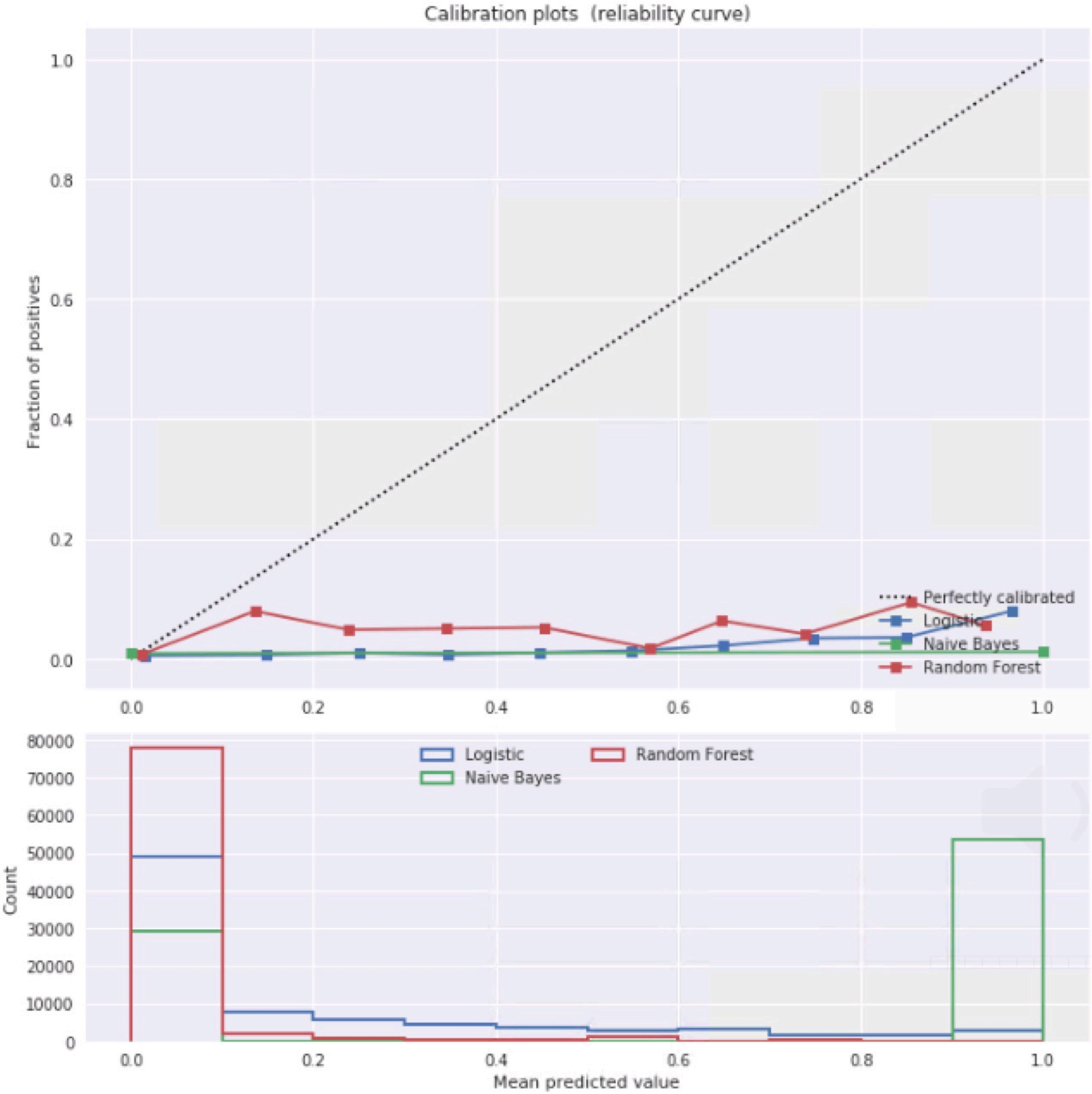

Calibration plots (reliability curve)
Fraction of positives
1.0
0.8
0.6
0.4
0.2
0.0
Perfectly calibrated
Logistic
Naive Bayes
Random Forest
0.0
0.2
0.4
0.6
0.8
1.0
Count
80000
70000
60000
50000
40000
30000
20000
10000
0
Logistic
Naive Bayes
Random Forest
Mean predicted value
